# Changes in cellular signaling patterns before and after stroke in the middle cerebral arteries of stroke prone spontaneously hypertensive rats

**DOI:** 10.1101/2020.09.11.292698

**Authors:** Killol Chokshi, Julie Warren, Krista Squires, Kayla A. Kitselman, Jules Doré, Noriko Daneshtalab

## Abstract

**Background:** Hemorrhagic stroke is associated with loss of middle cerebral artery (MCA) autoregulation in the stroke-prone spontaneously hypertensive rat (SHRsp). The signaling mechanism associated with the functional loss has yet to be defined. We hypothesize that physiological alterations coincide with changes to cerebrovascular inflammatory and contractile signaling and altered calcium signaling. METHODS: SHRsp rats were fed a high salt (4% NaCl) diet and sacrificed at 9 weeks of age for pre-stroke and after evidence of stroke for post-stroke samples. The MCAs were isolated for measuring protein levels using immunofluorescence (IF) & western blot (WB) for inflammatory signaling and contractile proteins. Tissues surrounding the MCA were analyzed for neuro-inflammation, neuronal damage, total and activated inflammatory proteins (ERK1/2 and p38MAPK), cerebrovascular contraction (PKC and MLC), and transient receptor potential V4 (TRPV4) expression. RESULTS: Our data show increase in activated inflammatory proteins after stroke with an associated decrease in expression of activated contractile proteins and TRPV4 channel expression compared to pre-stroke MCA. The post-stroke samples also show significant increase in neuro-inflammation and neuronal damage compared to pre-stroke samples.

**CONCLUSION:** An increase in activated/total (p38 MAPK &ERK1/2) is accompanied by a decrease in activated/total PKC & TRPV4 channel expression in post-stroke SHRsps. The decrease in vessel structural integrity and altered vascular tone of the MCAs may affect its ability to contract in response to pressure. Significant neuro-inflammation and neuronal damage in the brain tissues surrounding the MCA in post-stroke samples suggest MCA dysfunction is accompanied with neuronal and neural damage during stroke.

## INTRODUCTION

Stroke is one of the major cardiovascular diseases and a leading cause of functional impairment (partial paralysis, lapse in motor-coordination & instability) in North America(1, 2). The incidence of hemorrhagic stroke is approximately 20% of all stroke types. It is associated with having a high mortality and morbidity in adults (3), with the 5-year mortality rate of 50% in patients older than 45 years(4). Intracerebral hemorrhage (ICH), a type of hemorrhagic stroke, encompasses 10% to 15% of all stroke(5). The only treatment option available for ICH is surgery, with long-term hospitalization and rehabilitation being required upon surgical interventions as patients often suffer varying degrees of neurological dysfunction(6). ICH can arise due to multiple factors such as bursting (aneurysm) of weakened vessels(7), structural changes in the artery due to high blood pressure(8), or underlying vascular dysfunction resulting from an imbalance in vasoconstrictor/vasodilator factors(9).

The middle cerebral artery (MCA) directly mediates the pulsatile component of systemic blood circulation, and is involved in maintaining constant neural blood flow by a process called autoregulation (10). During chronic hypertension, consistent high blood pressure, above the MCA’s autoregulatory capacity, produces a 300-400% greater than normal increase in cerebral blood flow, known as autoregulatory breakthrough(11). This increase in flow is a result of loss of MCA vascular tone and forced dilation (12) due to impaired cerebral autoregulation(13). The loss of autoregulation is also associated with the inability of the MCAs to undergo pressure-dependent constriction (PDC), particularly in the stroke prone animal models(14). It is believed that the loss of MCA PDC advances cerebral perfusion and facilitates the development of ICH(15). Vascular dysfunction and inflammation have been implicated in many diseases, such as hypertension and stroke (16-18) and may be involved in the underlying signalling changes taking place within the MCA as stroke develops and PDC is lost.

Previous studies on isolated post-stroke MCAs have shown diminished response to PKC agonists, as well as loss of NOS activity, indicating functional deterioration of both the endothelial and vascular smooth muscle functions in the MCA(19). However, the underlying mechanism affecting the functioning of the vessel during stroke has not been investigated. The current work helps to identify the potential changes in protein expression occurring in the MCA before and after stroke in the SHRsp animal model. The study also investigates the extent of neuro-inflammation and neuronal damage in the brain tissue surrounding the MCA in pre-stroke and post-stroke SHRsps. The determination of cellular signalling patterns in MCA will help understand the vascular progression of stroke and its accompanying changes, leading to potential treatment options.

## MATERIALS AND METHODS

### Animals

All experimental procedures and animal breeding were carried out in compliance with the guidelines and recommendations, set forth and approved by the Animal Care ethics committee (Protocol Number: 15-30 ND), and the Canadian Council of Animal Care (Guide to care and Use of Experimental Animals, Vol 1, 2^nd^ Edition). They were performed at the Animal Care Facility, situated in the Health Science Center of the Memorial University of Newfoundland and Labrador. All efforts were made to minimize suffering during the experimentation and at the time of sacrifice and sampling. During the experimentation, stroke prone spontaneously hypertensive male rats (SHRsp; Charles River Laboratories, Quebec, Canada) were housed two per cage and bred in-house in ventilated cages under standard light cycle (12 hour light followed by 12 hour dark), controlled humidity, and temperature condition. The SHRsps used in the study were fed Japanese style stroke-prone high salt diet containing 4% NaCl (Zeigler Bros., Inc. Gardners, PA 17324 USA) from 5 weeks of age. Ad libitum access to food and water was permitted. At the time of sacrifice and sampling, the animals were anaesthetized with intraperitoneal injection of 50mg:10 mg/kg of ketamine:xylazine (Ketamine: Ketalean, Bimeda MIC, Animal Health Inc., ON, CAN & Xylazine: Rompun, Bayer Inc., ON, CAN).

### Experimental Design

The rats were divided into two experimental groups and sampled based on the previously established timeline of stroke(20). SHRsp rats were sacrificed before 10 weeks of age (mean of 9.3 weeks of age) for obtaining pre-stroke samples (n=24). To obtain the post-stroke samples, rats were monitored for external signs of behavioural distress and stroke-like symptoms. This consisted of weakness, lethargy, non-responsive to stimuli, pilo-erection, redness around eye, hunched back, sluggish movement, twitching, immobility & full and continuous seizures. The appearance of any and/or a combination of the mentioned distress signs were the determining factors for post-stroke samples (occurring approximately 15 weeks of age; n=24)(20). Samples of middle cerebral artery (MCA) and surrounding neural tissue were collected to perform immunofluorescence or Western Blot.

### Sample isolation and tissue processing

At the time of sampling, the animals were sacrificed according to methods listed above, and then exsanguinated by cardiac puncture in the left ventricle, using a heparinized 10 mL syringe and 18G needle. The brain was removed and placed in 1X Phosphate Buffered Saline (PBS) for isolating MCA and neural tissue. For western blot experiments, both MCA’s were dissected from the brain without stretching, flash frozen in liquid nitrogen and stored until analysis at −80° C. For immunofluorescence analysis, MCAs were cut alongside surrounding brain tissue, placed in a chip with embedding medium (Tissue Tek; Sakura Finetech Inc., CA, USA) and, flash frozen using liquid nitrogen prior to storage at −80° C. The remaining brain tissue was fixed in 10% neutral buffered formalin and embedded in paraffin for histological examination of neurological damage.

### Protein detection and analysis

MCA Samples were lysed in radioimmunoprecipitation assay (RIPA) Buffer within precellys beater tubes (Bertin Corp, MD, USA). Protein concentrations of each sample were determined by Bicinchoninic acid (BCA) assay using a standard curve generated with BSA standard (Pierce/ThermoFisher Scientific; IL, USA). Aliquots of equivalent total protein were separated by polyacrylamide gel electrophoresis and transferred to polyvinylidene difluoride (EMD, Millipore; Billerica, MA) membranes. Membranes were blocked in 5% non-fat dry milk in TBST prior to antibody detection of each specific protein of interest by incubating in blocking solution overnight on a rocking platform at 4C. Primary antibodies used were from Cell Signalling Technologies (New England BioLabs, ON, CAN). They included phosphorylated and total extracellular signal-regulated kinase (P-ERK 1/2, T-ERK 1/2), phosphorylated and total mitogen-activated protein kinase (P-p38MAPK, T-p38MAPK), phosphorylated protein kinase C (P-PKC), elastin microfibril interfacer 1 [EMILIN-1) and glyceraldehyde 3-phosphate dehydrogenase (GAPDH) Antibody for total PKC was from Santa Cruz Biotechnology (TX, USA; T-PKC).

Primary antibody binding was detected using a goat anti-rabbit IgG-Horseradish peroxidase secondary antibody (Santa Cruz Biotechnology: TX, USA), visualized with Supersignal West Pico Chemiluminscent Substrate (Pierce/ThermoScientific; IL, USA) by using the enhanced chemiluminescence imager: LAS400 (GE Healthcare, IL, USA). Membranes were then stripped and reprobed for quantitation of the corresponding total protein or GAPDH to allow for normalization of values based on total protein loading. Densitometric analysis of imaged blots was performed using Imagequant TL (GE Healthcare, IL, USA). For obtaining relative densitometry, ratio of phosphorylated over total protein and/or total protein over GAPDH was performed using Microsoft Excel 2010 (Microsoft Corporation, WA, USA).

### Tissue immunofluorescence

Flash frozen MCAs with surrounding brain tissue were sliced to 8 μm sections using a cryotome (Fisher Scientific, Pittsburgh, PA, USA), placed on charged slides (2-4 slices/slide), and stored at −20° C until processed for immunofluorescence (IF) studies. The expression of the following proteins were tested: phosphorylated and total myosin light chain [Cell Signalling Technologies; P-MLC (1:75); T-MLC (1:250)], transient receptor potential channel vanilloid 4 [TRPV4 (1: 100); Abcam, MA, USA] and filamentous and globular actin (F-Actin and G-Actin). On the day of testing, slides were thawed, washed in 1 x PBS. Samples staining for T-MLC required 0.5% SDS for antigen retrieval for 4 min at room temperature prior to tissue permealization and blocking. The blocking solutions consisted of either 2.5% Bovine Serum Albumin + 0.1% Triton-X in 1 X PBS (for P-MCL and T-MLC), or 10% normal Goat (for TRPV4) or Horse (for F- and G actin) Serum with 0.1% Triton-X in 1X PBS, incubated for 1 hr at room temperature. Following wash steps, the samples were incubated with the primary antibodies for the proteins of interest at concentrations indicated above, or with Alexa Fluor 488 phalloidin (ThermoFisher Scientific, ON, CAN) and Alexa Fluor 594 DNase I conjugate (ThermoFisher Scientific, ON, CAN) to stain F and G-actin respectively. On the second day, after 4 washes, sections were incubated in secondary antibody [Cy5 (1:400) with P-MLC; Cy2 (1:400) for T-MLC and Cy5 (1:300) for TRPV4; all Cy conjugated antibodies were from Jackson Immunoresearch, PA, USA]. 4’6-diamindino-2-phenylindole [DAPI (1:1,000); ThermoFisher Scientific, ON, CAN] was used as a nuclear counterstain.

In order to map potential areas of inflammation, sections were cut of brain tissue adjacent to the MCA and were stained for astrocytes and microglia/macrophages using GFAP-Cy3 (1:1,000; Sigma Aldrich, ON, CAN) and ionized calcium binding adaptor molecule 1[Iba-1; 1:1,000, Wako Chemicals, VA, USA]. Secondary antibody of Cy2 Goat Anti-rabbit (1:200; Jackson Immunoresearch, PA, USA) and DAPI (1:1,000) were applied in the second day according to established protocol (21).

For all the immunofluorescence staining, stacks of images at 1μm increments (a total of six slices) were collected for a Z-stack using confocal microscopy (FV1000; Olympus) with FluoView (Olympus) software. To perform semi-quantitative analysis, the images were converted to grayscale and analyzed using ImageJ software (U. S. National Institutes of Health). The vessel area (V) was determined and the mean gray value (MGV) of the vessel was measured. Mean gray value was determined to be the total pixel intensity of the measured area divided by the total measured area. Three random readings from the image background (B1, B2 & B3) were obtained. The MGV of the vessel was subtracted from the MGV of the background to obtain the actual optical fluorescence density of the vessel (M2). Thus, fluorescence intensity for the protein of interest was calculated as: M2=MGV (V) – MGV [(B1+B2+B3)/3]

### Quantification of neural damage

Neuronal and brain damage were analyzed using 5μm thick sections of hemotoxylin and eosin (H&E) stained cortical tissue by assessing cell vacuolation, neuron degeneration, areas of edema, and area of cell infiltration. The extent of neural damage was determined by a semi-quantitative scoring scheme of combining the scores obtained from four independent assessments outlined in Randell *et al*. (21)and adapted from Fedchenko *et al*.(22). In short, the grading scheme for cell death are ranked from 0 to 10, where cell count is graded in point system. The vacuolation is graded in factor of 10 cells per point, with the maximum number of vacuolation quantifiable being 80. The degenerating neurons were graded as 2 cells per point. Areas of edema and areas of cell infiltration (also indicators of brain injury and damage) were quantified separately, and graded by dividing each field into 10 sections, allotting 1 point for each section. The grading scheme for the areas is a modified version adopted from the “quickscore” system(23). A total final score was determined via summation of the four separate parameters to indicate total damage evident in the samples, as described by Randell *et al*.(21).

### Statistical analysis

Statistical analysis was performed using Excel 2010 (Microsoft Corporation, WA, USA) and SigmaPlot 12.5 (Systat Software Inc., CA, USA). Data were analyzed using unpaired two tailed student T-test. Value of p < 0.05 was considered statistically significant.

## RESULTS

### Downregulation of contractile signalling in the MCA after stroke

Changes in contractile signalling pathways were identified by performing western blot on the lysed sample of MCA obtained before (pre-) and after (post-) stroke. The ratio of phosphorylated PKC over total PKC (Figure 1B) was obtained for the immunoblots (Figure 1A), demonstrating a significant reduction in the ratio of active vs total PKD in post-stroke samples compared to pre-stroke samples (P<0.05). This corroborated our previous findings using PKC inhibitors indicating PKC vascular dysfunction in the MCA during stroke(19). No statistical difference was observed in the ratio of total PKC relative to GAPDH (loading groups) between sample groups (results not shown) demonstrating that there was no change in PKC expression due to stroke.

**Figure 1:**
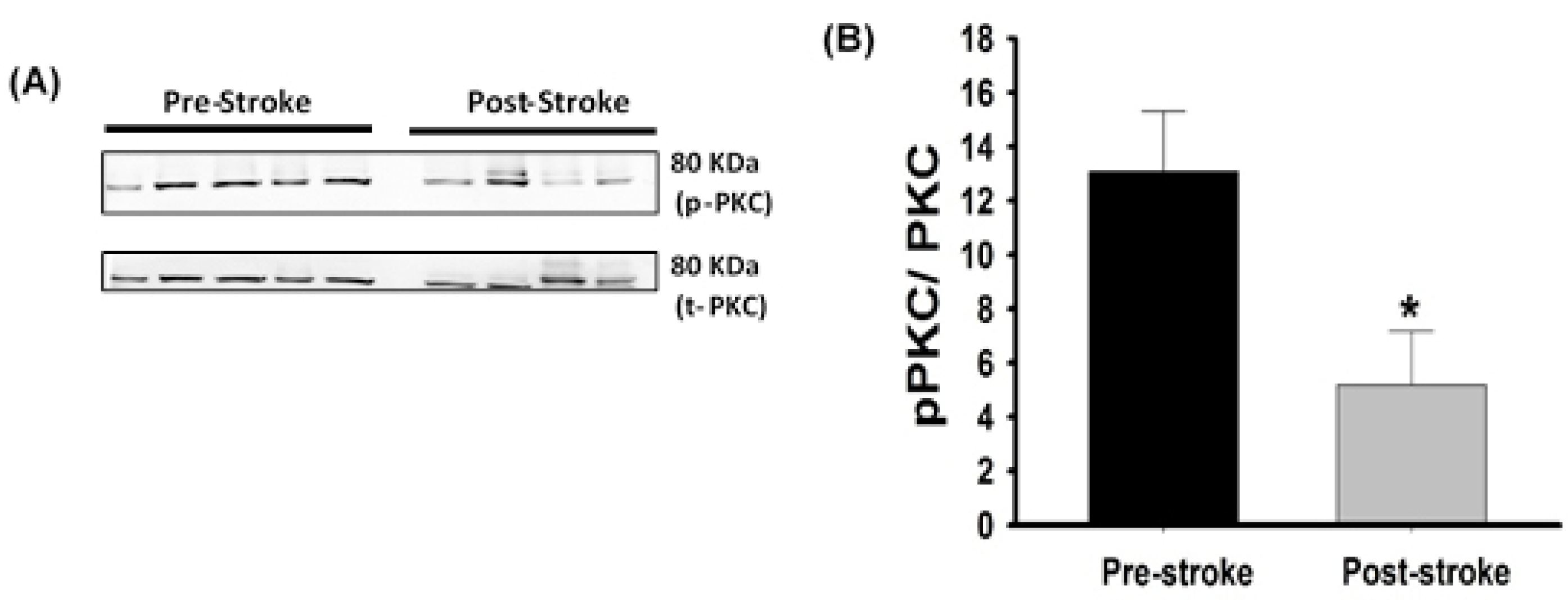
Phosphorylated & Total PKC levels in MCS pre and post stroke. **A)** Immunoblots showing the MCA levels of P-PKC with the corresponding immunoblot for total PKC to demonstrate loading controls. (p-PKC & t-PKC); **(B)** Relative densitometry for Phospho/Total PKC from MCA samples (n= 5) with * indicates p<0.05 analyzed using unpaired Student’s T-test.

Changes in the levels of myosin light chain (MLC), a vital signalling protein in contraction of the MCA, was analysed using immunofluorescence. The intensity of fluorescent signal from phosphorylated MLC stain was relatively low in both experimental groups resulting in a low dynamic range for analysis by mean gray value. Both total and phosphorylated MLC were detected primarily in the smooth muscle cells compared to the endothelium (Figure 2A, 2C). Semi-quantitative analysis showed significantly higher levels of phosphorylated MLC in post-stroke MCA’s compared to pre-stroke (Figure 2B; P<0.05). As we found with PKC, there was no significant difference between the mean gray value of total MLC in MCA of pre-stroke and post-stroke SHRsp rats (Figure 2D) demonstrating that there is not a decrease in the total PKC protein expression, but changes in activation levels occurring following stroke.

**Figure 2:**
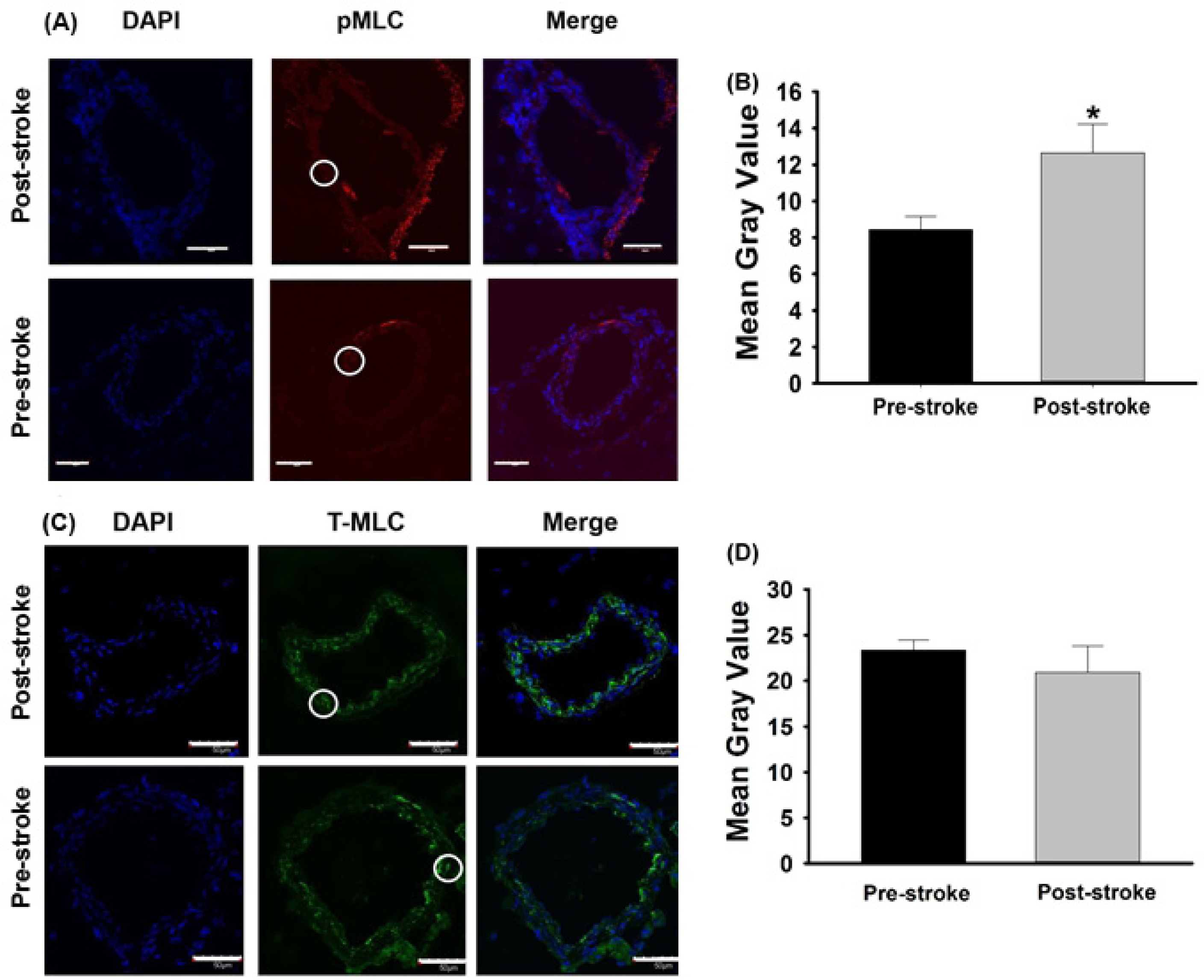
Representative images of MCA identifying MLC. A) Composite confocal Z-stacked images localizing Phospho-MLC2 within pre and post stroke MCA samples (P-MLC2 & DAPI) and; **(B)** Quantification of Mean Gray Value for phosphorylated MLC in MCAs of pre-stroke and post-stroke animals (n=6 per group). Representative images for **(C)** Total MLC Staining (T-MLC2 & DAPI) and; **(D)** Quantification of Mean Gray Value for total MLC in MCAs of pre-stroke and post-stroke animals (n=5 per group). * indicates p<0.05 using unpaired Student’s T-test. White arrows indicate phosphorylated and total MLC staining in the smooth muscle cells of MCA.

Since the formation of actin cytoskeletal structures are necessary for contractile machinery to function, we assessed for changes in the levels of F-Actin to G-Actin was measured in the MCA of pre-stroke and post-stroke by immunofluorescent staining (Figure 3A). Determination of the ratio of F:G-Actin using mean integrated intensity determination indicate a statistically lower ratio of F:G actin in MCA of post-stroke compared to pre-stroke samples (Figure 3B), indicating a loss in the actin structural components post-stroke.

**Figure 3:**
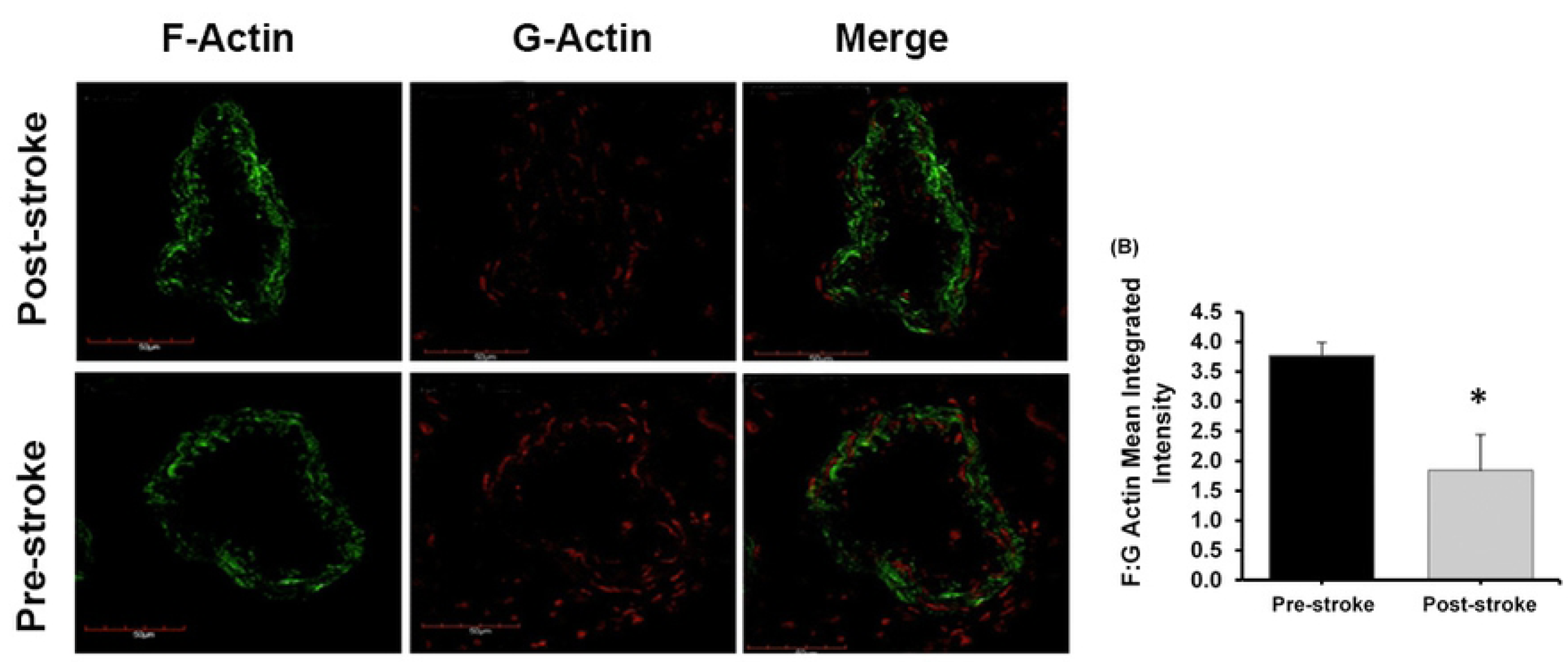
(A) Representative images for F-Actin and G-Actin obtained using confocal microscopy for both post-stroke and pre-stroke middle cerebral arteries. F-actin is represented in green and G-actin is represented in red. A composite of both is shown as ratio of F/G-Actin. **(B)** IF analysis of mean gray value of F/G-Actin in MCAs of pre-stroke and post-stroke animals. * indicates p<0.05 analyzed using unpaired Student’s T-test.

To further evaluate the structural integrity of the MCA vessels, we described the levels of EMILIN-1 in the MCA in order to characterize changes to the structural extracellular matrix of the vessel (ECM). Western blot analysis of the ratio of EMILIN-1 over GAPDH in the MCAs samples showed a significant decrease in the levels of EMILIN-1in post-stroke samples (Figure 4), furthering the hypothesis that MCA structural integrity is compromised following stroke.

**Figure 4:**
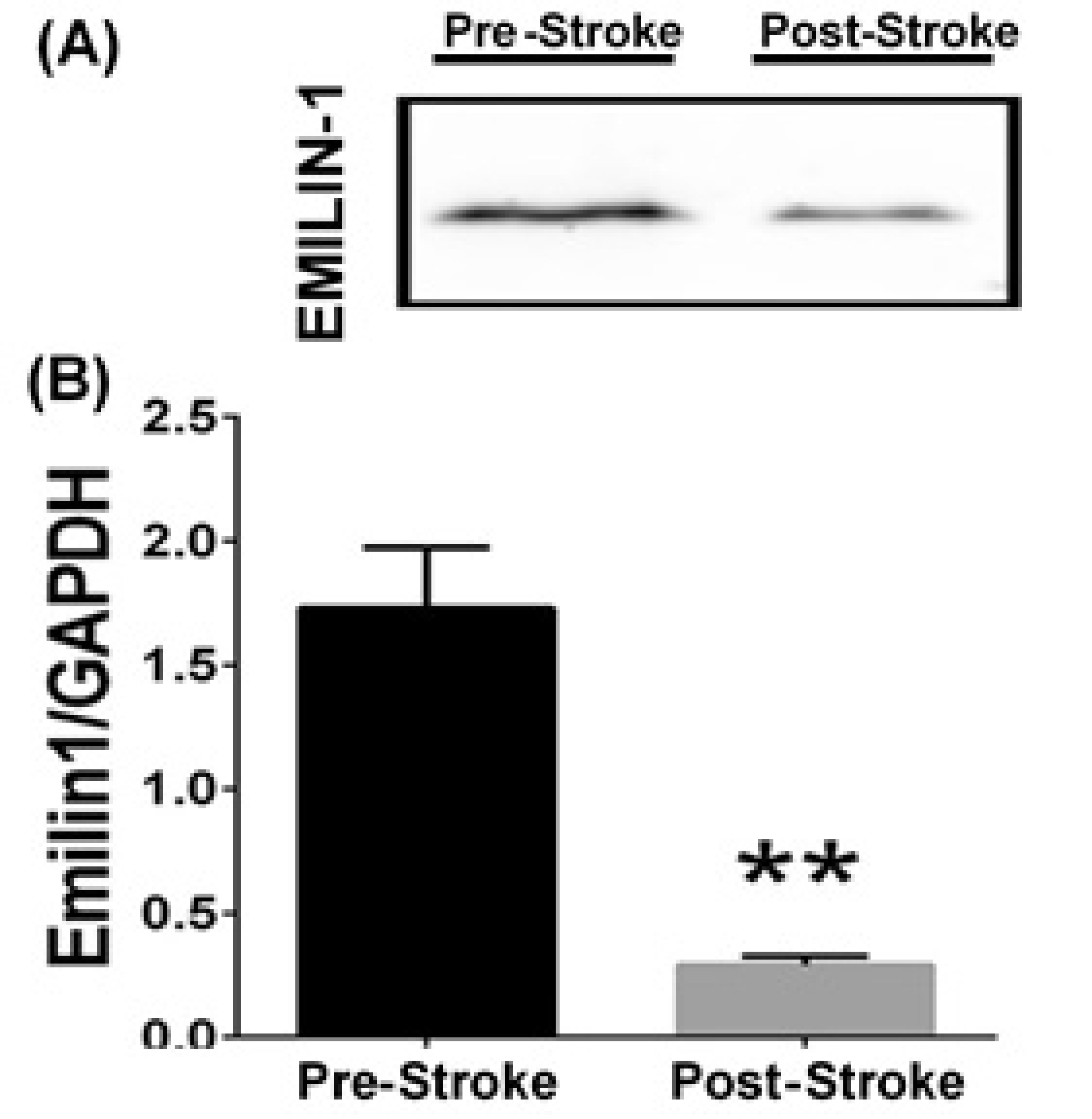
(A) Representative image for western blot bands for EMILIN-1 for both pre-stroke and post-stroke groups. **(B)** The graph shows ratio of EMILIN-1/GAPDH for pre-stroke and post-stroke groups (n=5). ** indicates p<0.001 analyzed using unpaired Student’s T-test.

### Downregulation of Calcium Channel expression after stroke

Thus far, we have demonstrated deficits in both the contractile signal transduction mechanisms and the contractile structural components in the MCA following stroke. We therefore wished to investigate calcium regulation within the MCA. MCA vessels from the pre-stroke and post-stroke SHRsp rats were immunostained to detect the expression of TRPV4 (calcium channel), often associated with vascular remodelling & stiffening in vessels(24). TRPV4 primarily localized in the endothelial layer compared to the smooth muscle cell layer of the MCA (Figure 5A). Semi-quantitative analysis of the mean gray value of TRPV4 staining showed a significantly lower expression of TRPV4 in post-stroke samples compared to pre-stroke samples (Figure 5B). PKC is known to be an important mediator in regulating activation of TRPV4 channels in endothelial cells (24,25), which may reflect the reason for lower TRPV4 channels accompanying lower activation of PKC (Figure 1) in post-stroke MCA.

**Figure 5:**
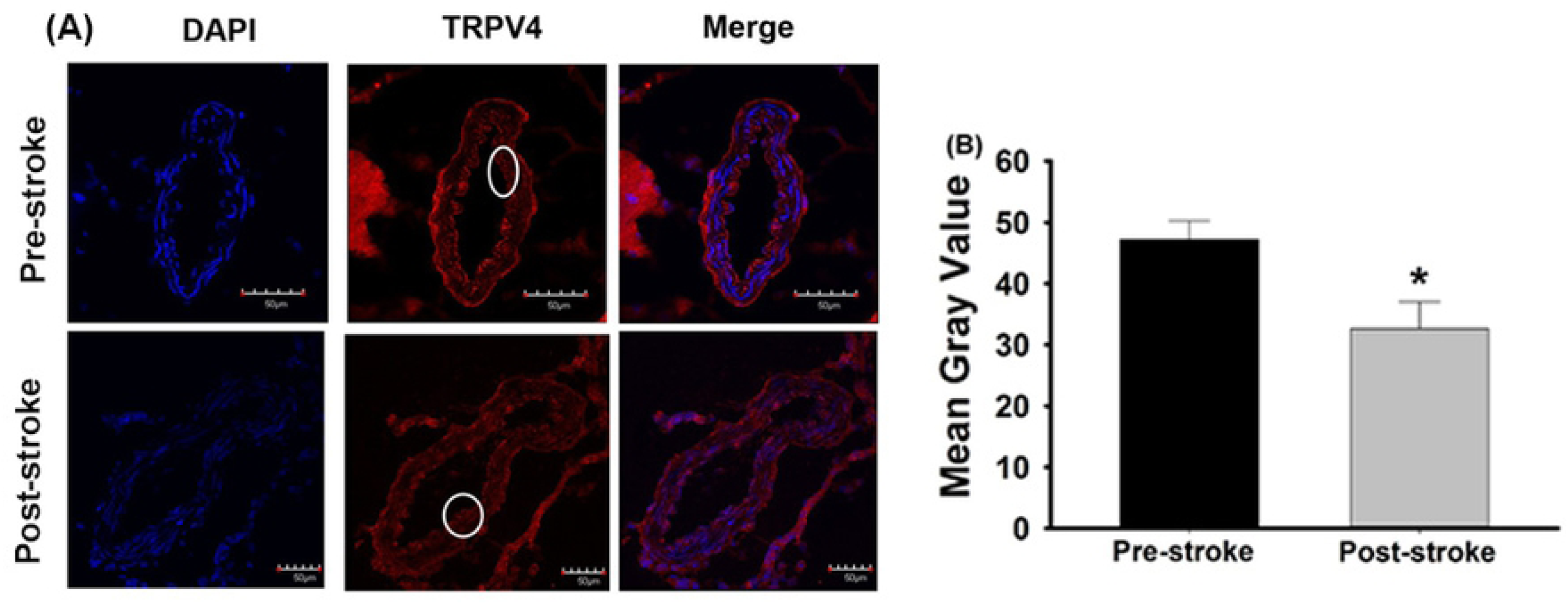
**(A) Representative images for TRPV4 Staining** (TRPV4 & DAPI). MCA were imaged as a Z-stack at 40x objective using confocal microscopy and fluoview software. Semi-Quantification of images performed by Image J software. **(B)** IF Analysis: Mean Gray Value for TRPV4 in MCAs of pre-stroke and post-stroke animals (n=6 per group). * indicates p<0.05 analyzed using unpaired Student’s T-test. TRPV4 staining has been indicated in the endothelium of MCA.

### Inflammatory activation in the MCA during stroke

We have shown that structural integrity and calcium signalling in the MCAs from post-stroke samples was decreased relative to pre-stroke samples. To address a potential underlying mechanism associated with changes to the structural integrity and calcium signaling in the MCAs, we evaluated the relative activation status of inflammatory mediators ERK and p38MAPK (Figure 6). Post-stroke samples show increased inflammatory activation in the MCA as the relative ratio for phosphorylated over total ERK1&2 was significantly higher in post-stroke samples compared to pre-stroke samples (Figure 6B; p <0.05). The increased activation of ERK in post-stroke samples was accompanied with increase in activated p38MAPK in the post-stroke samples compared to pre-stroke samples (Figure 6D). Together these findings suggest the presence of inflammatory damage in the MCA during stroke. There was no difference however for total ERK and total p38MPAK relative to GAPDH in either of the test groups indicating no change in total protein expression (results not shown).

**Figure 6:**
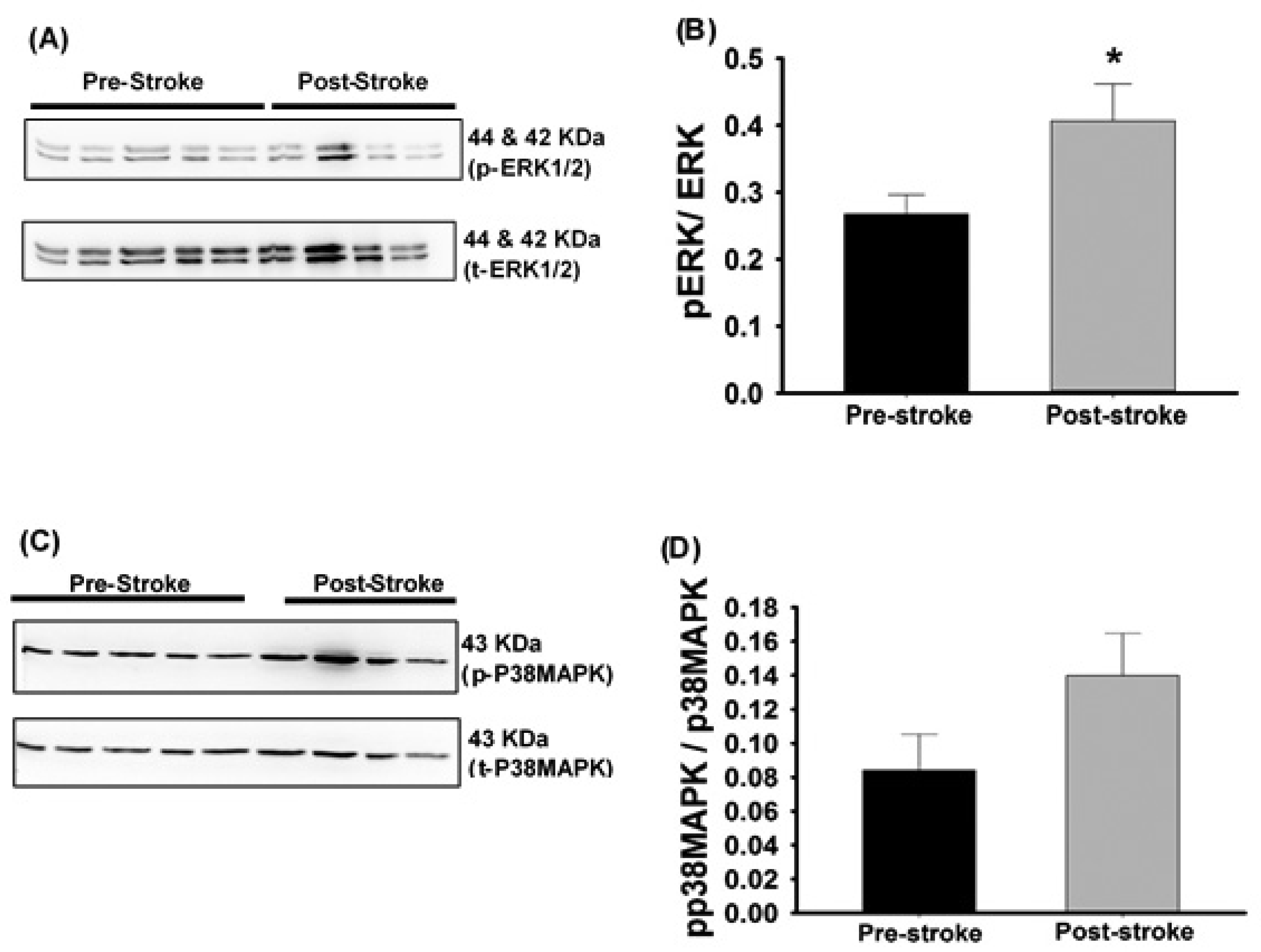
**(A) Representative image for Phosphorylated & Total P38MAPK bands** (P-p38MAPK (1:1000) & T-p38MAPK (1:1000)) and; **(B)** WB Analysis: Relative densitometry for Phosphorylated/Total P38MAPK. **(C)** Representative image for Phosphorylated & Total ERK bands (P-ERK1/2 (1:1000) & T-ERK1/2(1:1000)) bands; **(D)** WB Analysis: Relative densitometry for Phosphorylated/Total ERK1/2. * indicates p<0.05 analyzed using unpaired Student’s T-test.

### Increase in neuro-inflammation after stroke

The activation of inflammatory signalling markers in the MCA suggested that the brain itself may have been exposed to inflammatory damage as well. Therefore, microglia and astrocytes were imaged using brain slices immuno-stained with DAPI (nuclear stain), Iba-1(for total microglia) and GFAP (for astrocyte), respectively (Figure 7A). Semi-quantitative analysis of the number of activated microglia demonstrated a significant increase in the post-stroke samples compared to pre-stroke samples(Figure 7B) alongside an increase in astrocyte spread in post-stroke samples compared to pre-stroke samples (Figure 7C), indicating the presence of neuro-inflammation immediately following a stroke. The astrocyte spread and density were significantly greater near the MCA in the post-stroke samples, signifying considerable damage to the brain tissue surrounding the MCA.

**Figure 7:**
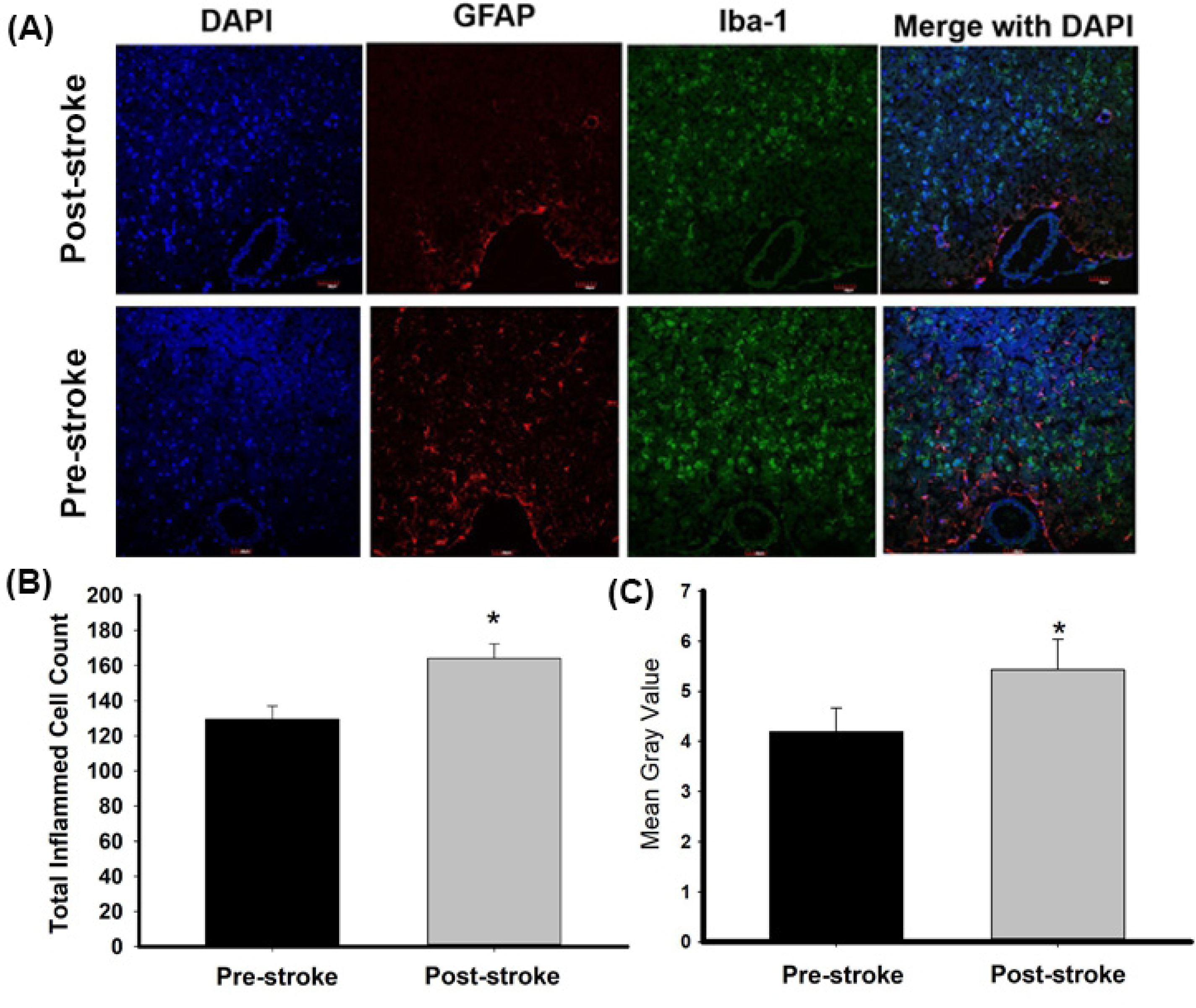
**(A) Representative images for Astrocytes & Microglia** (GFAP-Cy3 (1:1000), Iba 1 (1:1000), Cy5 (1;150) & DAPI (1:1000)). MCA and brain tissue sliced at 8μm and imaged at 20x objective using confocal microscopy and fluoview software. Semi-Quantification of imaged performed by Image J software. IF Analysis: **(B)** Mean Gray Value for Astrocytes & **(C)** Total Cell count of activated Microglia near middle cerebral arteries of pre-stroke and post-stroke animals (n=8 per group). * indicates p<0.05 analyzed using unpaired student’s t-test.

### Increase in neural damage near MCA during stroke

The brain slices from the pre-stroke and post-stroke SHRsp rats were stained using H&E staining and imaged to determine degrees of intracerebral and neural damage, particularly in the cortex in proximity to the middle cerebral artery (Figures 8A, B). The following four parameters were evaluated to determine the extent of damage in the brain during stroke using a semi-quantifiable scoring system: cell vacuolation, neuron degeneration, areas of edema, and cell infiltration. Total brain damage scoring indicated the post-stroke samples to have significant neural damage compared to pre-stroke samples (Figure 8C). Out of four parameters, degenerating neurons and cell vacuolation were the two most prominent changes observed in the post-stroke samples, indicative of greater neural damage in the brain regions near the MCA during stroke.

**Figure 8:**
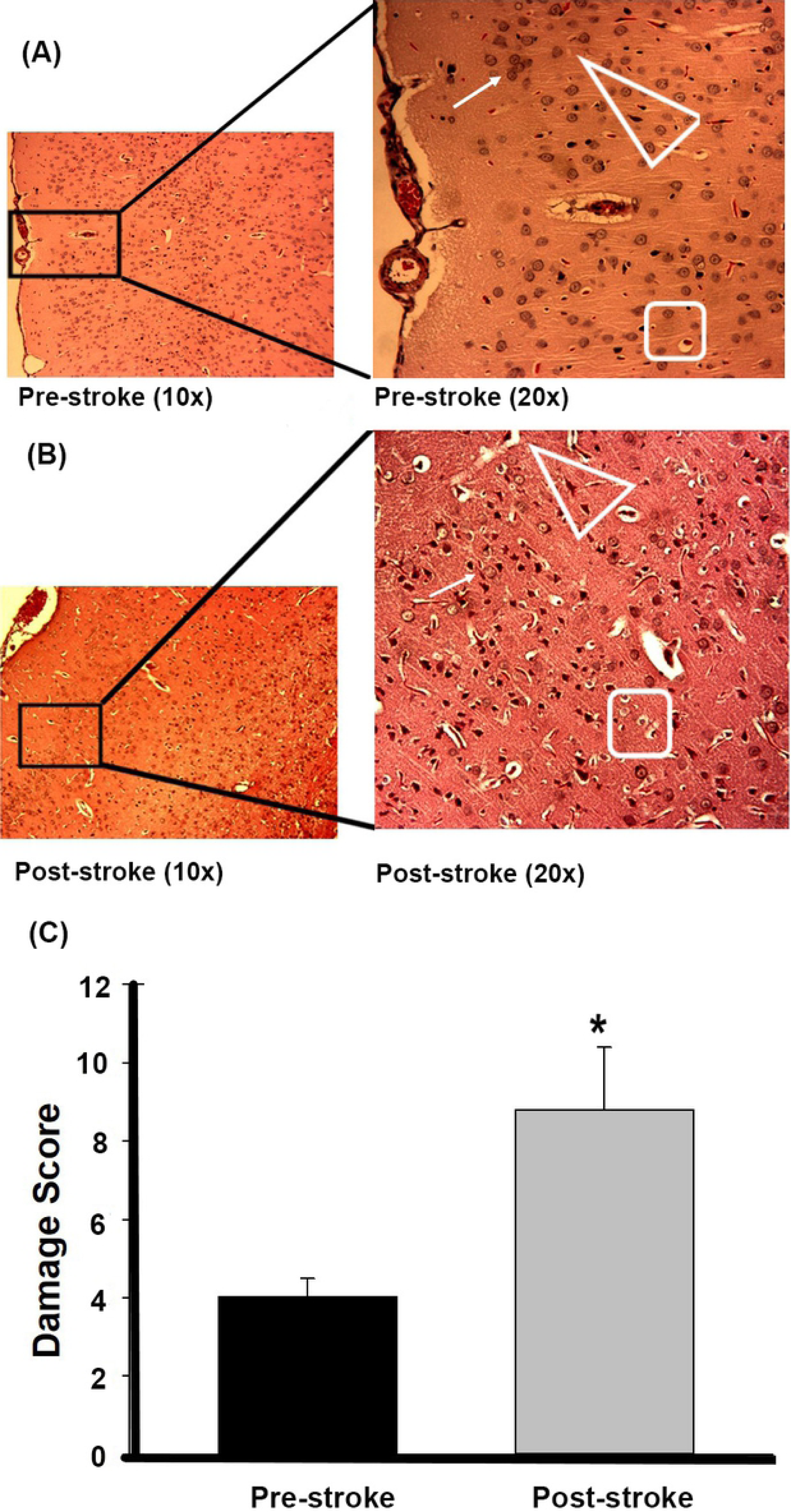
Brain pathology using H&E stains of brain slices (6 μm) imaged at 10× and 20× objectives for pre-stroke **(A)** & post-stroke **(B)**. Areas of neural cell vacuolation (white rhombus), neural degeneration (white triangle), and cell infiltration of inflammatory origin (long white arrow) are indicated between the images **(A–B)**. Total Neural Damage Scoring by H and E staining of brains, for pre-stroke (n=6) and post-stroke (n=4) groups. * indicates p<0.05 analyzed using unpaired Student’s t-test.

## DISCUSSION

The stroke-prone spontaneously hypertensive rat (SHRsp) emulates physiological aspects of spontaneous hemorrhagic stroke seen in humans(20). When fed a 4% NaCl diet they develop high blood pressure at approximately eight weeks of age, followed by spontaneous hemorrhagic stroke at 11-15 weeks with, limb weakness, motor uncoordination, weight loss, dehydration, twitching and seizures(20, 25). Previous research by our group found significant pathophysiological differences in the vascular function of the MCA between pre-stroke and post-stroke SHRsp, with a loss of autoregulation and associated ability to undergo pressure dependent constriction in the MCAs in the event of stroke(14, 26). In this investigation, we determined the cellular signalling changes associated with contractile and inflammatory mechanisms in the MCA, potentially contributing to occurrence of intracerebral hemorrhage.

The MCA undergoes both vasodilation and vasoconstriction to maintain the consistent blood flow, with the vascular smooth muscle cells and endothelium being the contributors in both of the processes. Numerous contractile stimuli, such as an increase in intraluminal pressure and intracellular calcium(27, 28) lead to smooth muscle cell contraction(29). The loss of autoregulation in cerebral arteries exhibited post-stroke is due to the loss of normal functioning of the vessels to respond to pressure and absence of the normal contractile cycle. Normally, increased calcium levels activate Myosin Light Chain Kinase (MLCK) to phosphorylate MLC, resulting in contraction of vascular smooth muscle(28). It is possible that changes in phosphorylation of MLC would affect contractility of the vessel. We found a significant increase in activated MLC in the post-stroke MCA, with no change in total MLC, suggesting upstream signalling events are stimulating the contraction process. However, activated MLC after stroke has a lower availability of F-actin to form cross-bridge cycling. Actin polymerization is necessary for actin-myosin cross-bridge cycling, being a fundamental mechanism for tension development and contraction in all muscles, including smooth muscle cells(30). Although the phosphorylation status of MLC is independent of actin polymerization, inhibition of F-actin polymerization either by pharmacological or molecular approaches results in attenuated smooth muscle contraction(31). F-Actin is vital for proper contraction of smooth muscle, as shown on isolated vascular smooth muscle cells and tracheal smooth muscle layers, where an increase in F-actin & decrease in G-Actin corresponds to contractile stimuli(30). The significant decrease in ratio of F/G-actin in post-stroke MCAs, despite higher levels of phospho-MLC, likely contributes to decrease in ability to maintain stable vascular contraction and may be the result of loss of vascular integrity in our animal model. As the development of contractile force in smooth muscle requires actin polymerization and phospho-MLC to work in parallel signalling pathways, absence of either pathway hinders the normal functioning of the smooth muscle. Studies have shown ERK to mediate vasoconstriction via activation of MLCK, leading to increased phosphorylation of MLC(32). Our data indicates an increase in activation of MLC alongside an increase in activated ERK, in post-stroke MCA’s, suggesting an inter-connected signalling pathway to compensate for chronic high blood pressure in order to attempt to rescue normal functioning post-stroke.

Contraction of VSMC involves activation of multiple interlinked pathways that include phosphorylation of MLC. Activation of PKC increases the myofilament force sensitivity to [Ca^2+^] and MLC phosphorylation, thus maintaining contraction(33). Studies in bovine MCAs have also shown that PKC increases vascular tone by decreasing myosin light chain phosphatase (MLCP) activity, thus increasing MLC phosphorylation(34). Our analysis of MCA’s for phospho-PKC (all isoforms; (35)) showed lower levels of phospho-PKC in post-stroke MCA. This finding corroborates our previous findings that MCAs had significantly reduced PKC activation capacity post-stroke when stimulated with phorbol ester(36). Intracellular calcium and DAG both regulate activation of various isoforms of PKC (37). The intracellular concentration of calcium triggers association of PKC isozymes with the membrane to allow DAG interaction with PKC to stimulate activity, as seen in studies in diabetic vasculature where DAG and its analogues activate specific isoforms of PKC(38, 39). Although no specific isoform of PKC has been correlated with MCA vascular tone, the down regulation of either DAG or calcium would decrease the activation of PKC in post-stroke samples, and thus a decrease vascular tone.

Interestingly, studies have shown upregulation of PKC signalling results in TRPV4-mediated [Ca ^2+^] influx in endothelial cells, via phosphorylation at the Ser824 site (40). In this study, there was a decrease in TRPV4 expression coincidentally with a decrease in active PKC post-stroke. TRPV4 is essential in the regulation of vascular function during mechanical stress(41). Our observed decrease in TRPV4 expression may be due to the decrease in PKC activation. There may also be a negative feedback loop in which decreased TRPV4 levels decrease calcium influx and PKC activity, which further decreases TRPV4 functionality and ultimately decreasing intracellular calcium levels and inhibiting contraction.

The decrease in TRPV4 expression is associated with the vascular remodelling and stiffening, accompanied by deposition of calcium and thinning of the endothelial layer (42), making the vessel less responsive to various stimuli. As TRPV4 channels are highly osmo-sensitive and mechano-sensitive (43), the absence or diminished detection of either stimuli due to thinning of the endothelial layer during vascular stiffening may impact the expression of TRPV4, as seen in the post-stroke samples.

The altered pathways we identified suggest inhibited contractile mechanisms during stroke may partially be responsible for the loss of pressure dependent constriction seen in the post-stroke samples. However, the underlying mechanism resulting in these changes is unknown. One likely candidate is inflammatory signalling in the MCA just prior to stroke. The pre-stroke samples we analyzed represent the steady state levels long before stroke while post–stroke samples represent the steady state immediately following stroke. MAPK cascade (p38 MAPK and ERK1/2) are important inflammatory signalling pathways controlling a broad spectrum of cellular processes and stress responses (18, 44). An increase in ERK signalling results in increases in inducible nitric oxide synthase, cyclooxygenase-2 (45), while p38 MAPK has been specifically implicated in endothelial injury (45). Activation of both p38 MAPK and ERK 1/2 upregulates transcription factors leading to generation of inflammatory mediators (IL-6, TNF-α and IL-1β; (46-48)). Additionally, p38 MAPK activated by TLR promotes production of pro-inflammatory cytokines, TNF-α, IL-1β, IL-6 and IFN-γ (45, 49). Our results indicate the activated p38 MAPK and ERK1/2 were significantly increased post stroke, indicating the presence of a pro-inflammatory mileu in the cerebral vasculature. This suggests a positive feedback loop with an increase in stress related signal causing more inflammation in the vessel in the post-stroke MCA.

Different molecular patterns associated with sensing functional or structural changes in the vasculature activate and increase ERK phosphorylation in the post-stroke MCA (49). It is also possible that the increase in activated ERK, post-stroke, is a compensatory mechanism for the loss of contractile function, as increase in ERK pathway has shown to activate actin assembly causing vessels to contract (50). The role of ERK in potentiating vascular dysfunction likely starts during the later pre-stroke stage. Exposure to chronic high blood pressure prior to stroke causes an increase in phosphorylated ERK1/2 levels in rat cerebral and coronary arteries (51). During exposure to chronic high blood pressure, the vessels suffer extreme shear stress and circumferential stretch, causing extensive endothelial damage, and extracellular matrix (ECM) imbalances (52), likely resulting in a strong inflammatory response (53). Evidence of MCA ECM remodeling is indicated by the observed decrease in EMILIN-1 levels. The decrease in EMILIN-1 in post-stroke MCAs would directly affect vascular ECM stability(54) and integrity.

MAPK activity is also linked with microglial activation due to increased inflammatory activity. In striatum of rats that had induced ICH, inhibition of MAPK pathways decreased survival of activated microglia (55) and phosphorylated p38 MAPK localizes to neurons and microglia of ischemic brain tissue in the hippocampus (56). The brain regions surrounding the post-stroke MCAs, the anterior region extending from insula with the opercular segments (parietal and temporal), showed significantly higher levels of activated microglia compared to the brain regions around the pre-stroke MCA. This reflects an increase in neural inflammation in tandem with increased vascular inflammation. Activated microglial cells have been shown to reduce proliferation, survival and function of new neurons (57), as they are known to produce pro-inflammatory cytokines TNF-α, IL-1β, IL-6 (58) and CXCL2 which promote neuro-inflammation and recruitment of leukocytes to the brain (59). Activated microglia induce active metalloproteinase-9 generation (60), known to be involved in vascular remodelling through degradation of extracellular matrix (ECM), as well as activating cytokines/chemokines in the brain and cerebro-vasculature (60).

In addition to the activation of microglia post-stroke, a significant increase in astrocyte spread in the post-stroke brain regions proximal to the MCA is seen. Increase in astrocyte accumulation near the site of damage is believed to contribute to local inflammatory responses by producing pro-inflammatory cytokines (61). In addition, Astrogliosis at the site of damage (62) contributes to degeneration of synapses and axonal neuro-filaments (63), leading to loss of neuronal conduction. The significant increase in astrocyte spread near the MCA region in post-stroke samples indicate the magnitude of damage after stroke. Astrocytes are also known to regulate vascular smooth muscle cells and support structural integrity in small cerebral vessels in deep brain regions (64). In adult mice, absence of astrocytic laminin led to impaired function of VSMC and vascular wall disassembly, making it an important mediator in structural integrity of cerebral vessels. It maybe that the increase in astrocytes near the MCAs are a compensatory mechanism to maintain some vascular integrity post-stroke. Alternatively, activation of astrocytes can induce accumulation of excessive glutamate during brain damage (65) causing damage in surrounding brain tissue. Analysis of the downstream signalling of astrocyte-activated proteins would further illuminate the specific stage of the inflammatory process in our model.

In order to clarify the effect of stroke on the neighbouring neural tissue we further analyzed changes in neural cell vacuolation, degeneration, area of edema and cellular infiltration of inflammatory origin, in brain regions proximal to the MCA. Neuron degeneration and vacuolation have been associated with cell death, and are associated with a broad range of inductive stimuli (66). Post-stroke brain regions near the MCA showed elevated levels of neural damage, relative to pre-stroke, with degenerating neurons and cell vacuolation as the most prominent differences observed, indicating substantial neural damage in the brain regions near the MCA following stroke. Axon degeneration has been widely seen in neurodegenerative diseases, including stroke and motor neuropathies (67), and is associated with impaired transmission across the neural network and cytoskeletal breakdown (67). This would cause a significant decrease in the neural network, making the brain regions following stroke less responsive and functional.

Our results showed significant differences in inflammatory and contractile signalling pathways before and after stroke, however the exact timeframe of damage is unknown. It is unclear if many of the signalling changes we have identified occur just prior to, or follow from the consequences of the stroke event and what other structural changes occur in the vessels to promote dysfunction. We demonstrate that the combination of increased inflammatory expression and decreased contractile signalling is responsible for the loss of auto-regulatory mechanisms in the MCA after stroke, and correlate with the functional results we have previously published. These changes during stroke are accompanied with an increase in neural damage as well as neuro-inflammation, affecting the brain region surrounding the MCA during stroke. We believe the presence of inflammation in the MCA and surrounding brain region induce the structural and functional changes seen in MCA after stroke and significantly contribute to the severity of ICHs. The use of specific anti-inflammatory and specific calcium channel blockers as potential treatment options for ICH would be important to consider, in light of the pathophysiology associated with ICH.

## Abbreviations

BCA: Bicinchoninic acid
COX-2: Cyclooxygenase-2
EMILIN1: Elastin microfibril interface–located protein 1
GAPDH: Glyceraldehyde 3-phosphate dehydrogenase
GFAP: Glial fibrillary acidic protein
HRP: Horseradish peroxidase
Iba-1: Ionized calcium binding adaptor molecule 1
iNOS: inducible nitric oxide synthase
MCA: Middle Cerebral Artery
MGV: Mean gray value
MLC: Myosin Light Chain
MMP-9: Matrix metalloproteinases-9
NBF: Neutral Buffered Formalin
NGS: Normal Goat Serum
PBS: Phosphate Buffered Saline
PDC: Pressure Dependent Constriction
PFA: Paraformaldehyde
PKC: Protein Kinase C
PVDF: Polyvinylidene fluoride
RIPA: Radioimmunoprecipitation assay
SDS: Sodium Dodecyl Sulfate
SHRsp: Stroke prone Spontaneously hypertensive rats
TBS: Tris-buffered saline
TBST: Tris-buffered Saline & Tween 20
TRPV4: Transient Receptor Potential Cation Channel

